# Reengineering the Specificity of the Highly Selective *Clostridium botulinum* Protease via Directed Evolution

**DOI:** 10.1101/2020.09.29.319145

**Authors:** Rebekah P. Dyer, Hariny M. Isoda, Gabriela S. Salcedo, Gaetano Speciale, Madison H. Fletcher, Linh Q. Le, Yi Liu, Shiazah Z. Malik, Edwin J. Vazquez-Cintron, Andrew C. Chu, David C. Rupp, Birgitte P.S. Jacky, Thu T.M. Nguyen, Lance E. Steward, Sudipta Majumdar, Amy D. Brideau-Andersen, Gregory A. Weiss

## Abstract

The botulinum neurotoxin serotype A (BoNT/A) cuts a single peptide bond in SNAP25, an activity used to treat a wide range of diseases. Reengineering the substrate specificity of BoNT/A’s protease domain (LC/A) could expand its therapeutic applications; however, LC/A’s extended substrate recognition (≈60 residues) challenges conventional approaches. We report a directed evolution method for retargeting LC/A’s substrate and retaining its exquisite specificity. The resultant eight-mutation LC/A (omLC/A) has improved cleavage specificity and catalytic efficiency (1300- and 120-fold, respectively) for SNAP23 versus SNAP25 compared to a previously reported LC/A variant. Importantly, the BoNT/A holotoxin equipped with omLC/A infiltrates neurons and retains its SNAP23 activity. The identification of substrate control loops outside BoNT/A’s active site could guide the design of improved BoNT proteases and inhibitors.

**One Sentence Summary:** Directed evolution of the BoNT/A protease targets a new cellular protein, SNAP23, expanding its therapeutic potential.

## Main Text

Therapeutic proteases must select a narrow range of substrates to ensure safety and efficacy (*1*). For example, BoNT/A hydrolyses a single peptide bond in its sole substrate SNAP25 (*2*). This cleavage of SNAP25 disrupts vesicle fusion, neurotransmitter release, and neuro-muscular communication. The remarkable specificity of BoNT/A and its months-long duration of clinical benefits allows its application to a broad and expanding list of neuronal indications, ranging from spasticity to facial wrinkles and migraines (*3*). The complete toxin, or holotoxin, has a heavy chain (HC/A) and light chain (LC/A), which cooperate to infiltrate peripheral neurons and cleave SNAP25, respectively. The HC/A binding domain enters the cell through receptor-mediated endocytosis (*4*, *5*). Upon endocytic escape mediated by the translocation domain within HC/A and subsequent disulfide reduction, LC/A recognizes and cleaves SNAP25 (*6*).

Retargeting the delivery or substrate selectivity of BoNT/A could expand its therapeutic uses. For example, antibody fragments (*7*, *8*), growth factors (*9*), lectins (*10*), or other proteins (*11*) fused to the HC/A translocation domain could successfully deliver BoNT subtypes to new cell types to potentially treat pain, neuroendocrine disorders, and cancers (*12*). Several non-neuronal SNARE proteins offer attractive therapeutic targets for reengineering the proteolytic specificity of LC/A. For instance, SNAP23 mediates the release of pro-inflammatory molecules (*13*, *14*), secretion of matrix metalloproteinases that promote tumor invasion (*15*), and hypersecretion of mucin in asthma (*16*). Though not a substrate for wild-type LC/A (wtLC/A), SNAP23 is the closest homolog to SNAP25, and SNAP29 is the next closest homolog (**Fig. 1A**). Thus, we sought to reengineer LC/A specificity for SNAP23 to expand therapeutic uses of BoNT/A biopharmaceuticals.

**Fig. 1.**
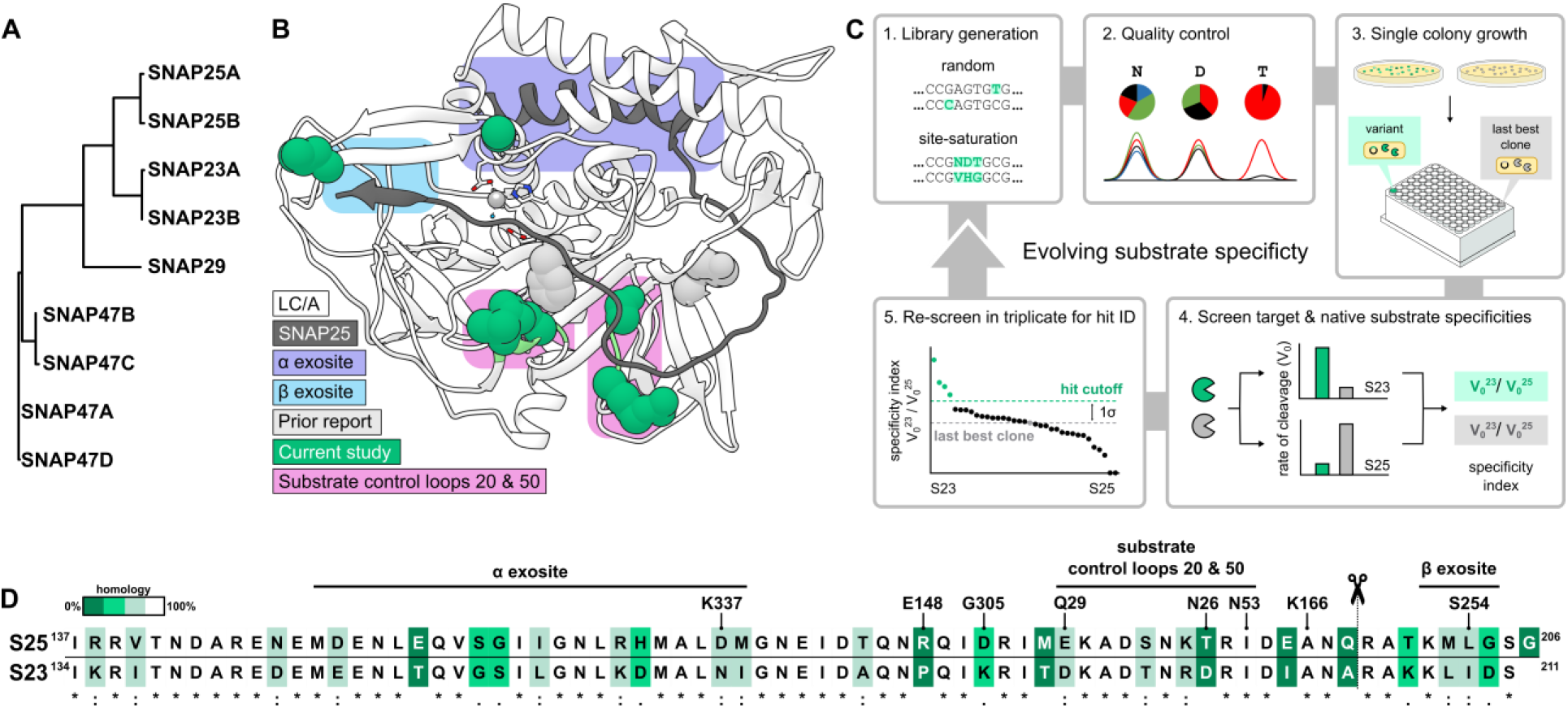
Directed evolution overview. (**A**) SNAP25 sub-family proteins and their isoforms (CLUSTAL 2.1 (*25*)). (**B**) Co-crystal structure (PDB: 1XTG, (*26*)) of LC/A (white) and SNAP25 (dark gray). Eight substitutions in LC/A drive SNAP23 specificity (teal) through substrate control loops (pink) alongside prior substitutions (light gray). (**C**) Platform for the directed evolution of SNAP23 substrate specificity. 1. Random or site-directed mutagenesis (e.g., site-saturation); 2. QC by DNA sequencing; 3. High-throughput protein production; 4. Measure V_0_^23^ and V_0_^25^ for substrate specificity; 5. Confirmation screens. The most specific and consistent variant from the DARET assay entered the next round of directed evolution. (**D**) Sequence alignment of substrates used for screening (UniProt: P60880, O00161). The SNAP binding exosites in LC/A (residue numbers above) and cleavage site (scissors) are shown. The gradient of color indicates homology from identical (white, *), to strongly similar (light gray,:), weakly similar (teal, .), or dissimilar (dark teal).

LC/A obtains its tremendous enzymatic specificity by interrogating ≈60 residues of its substrate (**Fig. 1B**). Conventional platforms for protease reengineering are limited to short recognition sequences (*17*–*19*). Therefore, progress towards evolution of LC/A to cleave SNAP23 has largely relied on *in silico* studies (*20*). For example, adding four computationally derived mutations (E148Y/K166F/S254A/G305D) to LC/A generates some SNAP23 cleavage activity (*21*, *22*). However, this quadruple mutant of LC/A (qmLC/A) remains highly specific for its native substrate, SNAP25 (**Fig. S1**). Here, directed evolution was applied to retarget qmLC/A to specifically cleave SNAP23 over SNAP25.

To screen for substrate specificity, depolarization after resonance energy transfer (DARET) (*23*, *24*) measured LC/A cleavage of SNAP23 and SNAP25. Here, >70 residues of each substrate were sandwiched between GFP and BFP (shown schematically in **Fig. S2**). This assay features exceptionally low background compared to a FRET-only assay and increased polarization upon proteolysis (*23*). During each round of directed evolution, normalizing the initial rate of proteolysis for SNAP23 (V_0_^23^) to the analogous rate for SNAP25 (V_0_^25^) yielded a substrate specificity index, which accommodated variation in enzyme concentration (**Fig. 1C**). The assay proved effective at revealing even small gains in substrate specificity.

Error-prone PCR of the gene encoding qmLC/A in Round 1 uncovered three variants with potentially improved SNAP23 specificity: N240S, E201D/D203V, and K364R/Y387N. The latter two variants exhibited inconsistent SNAP23 specificities in triplicate follow-up screens, perhaps due to protein insolubility. Only N240S qmLC/A yielded a consistent increase in SNAP23 specificity compared to qmLC/A. Additional site-saturation at this position in Round 2 identified N240A as improving SNAP23 specificity approximately 1.3-fold or >1σ above the mean specificity for qmLC/A data at the same screening dilution (σ = standard deviation) (**Fig. 2A**). This residue exists in a semi-ordered loop approximately 6 Å away from the beta hairpin of the beta exosite, an important contributor to binding SNAP25.

**Fig. 2.**
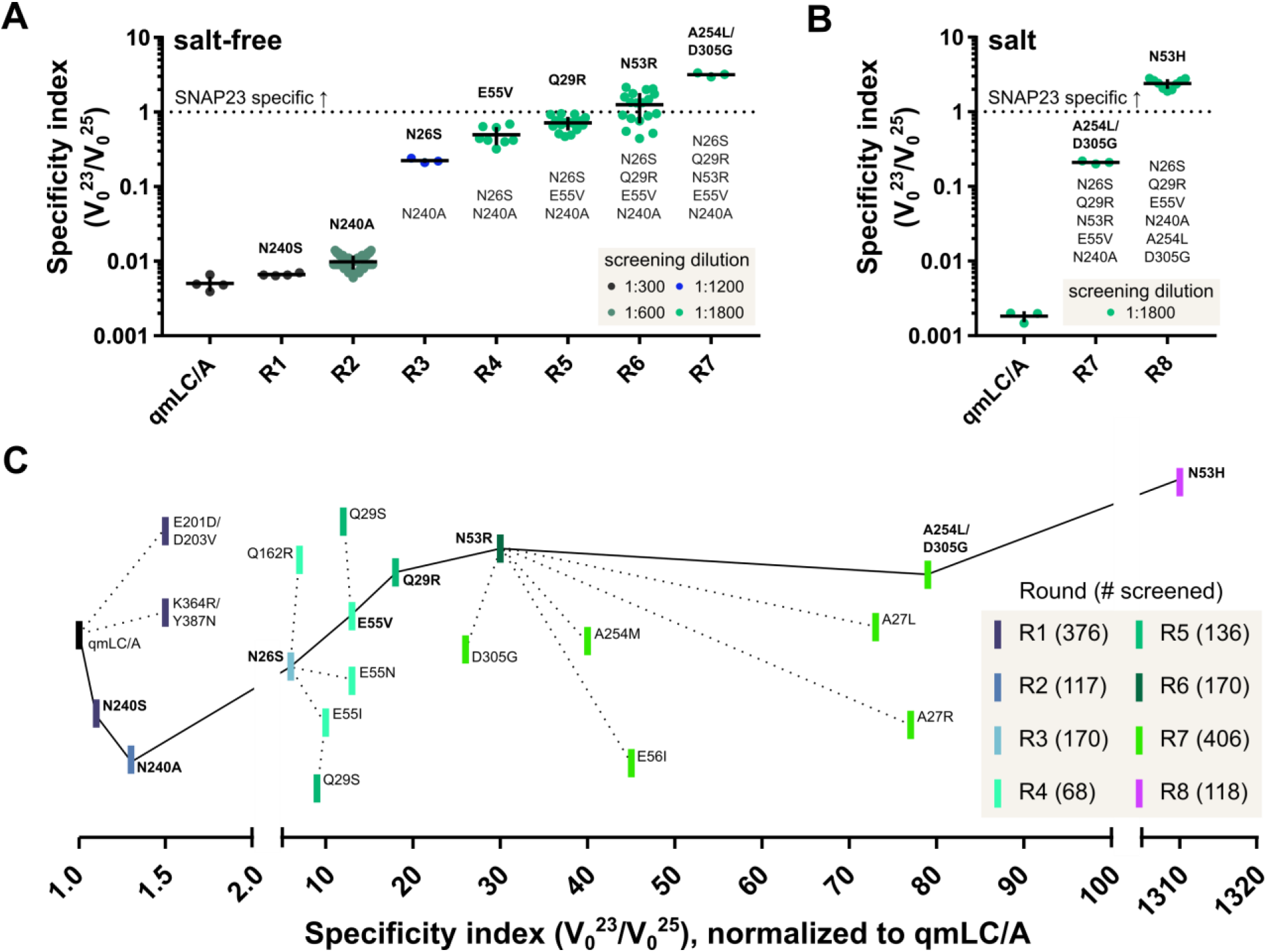
Directed evolution of a SNAP23-specific LC/A. (**A**) Improvement in specificity index through directed evolution in salt-free buffer. The most specific variant from each round was assayed multiple times (replicates shown). The average specificity index (horizontal bars) and standard deviation (error bars) are shown. The final dilution of cell lysates for screening is indicated by color code. (**B**) Increased specificity index achieved through directed evolution in salt buffer. (**C**) Roadmap for directed evolution showing the fold increase in specificity indices over eight rounds (R1 to R8). As shown in **Fig. S1**, the substrate specificity of the qmLC/A variant is sensitive to screening dilution; therefore, each point represents the variant’s average specificity normalized to the qmLC/A average specificity at the corresponding screening dilution. Advancement to the next round (solid line) weighed specificity index, protein solubility, and stability.

Subsequent rounds of directed evolution were guided by rational design, including inspection of the LC/A-SNAP25 co-crystal structure (**Fig. S3**) (*26*) and prior work (*21*, *22*). Rounds 3 to 7, for example, included residues with high B-factors, which can be productive sites for substitution (*27*). Investigating these sites with an iterative mutagenesis method (*28*) improved specificity for SNAP23 up to 80-fold over qmLC/A in the DARET screens (**Fig. 2A**). In each round, Q_pool_ analysis (*29*) was used to assess library quality. The TopLib program estimated the numbers of variants required for screening in each round (*30*).

Later rounds of directed evolution identified a sensitivity to the screening and assay conditions. In early rounds, variants were diluted and screened in a “salt-free” assay buffer (50 mM HEPES pH 7.4, 0.05% Tween with no added salt) chosen to maximize LC/A cleavage activity *in vitro* (*23*). These screening conditions yielded LC/A variants that cleaved SNAP23 at over three times the rate of SNAP25 cleavage (**Fig. 2A**). However, the addition of salt reversed these gains in SNAP23 specificity for the selectant from Round 7 (**Fig. 2B**, **Fig. S4**). The results vividly illustrate the First Law of Directed Evolution, “you get what you screen for” (*31*), as screening took place in salt-free buffer.

Since we sought protease variants having specificity for SNAP23 within the salty environment of cells, we addressed this issue with a final round of mutagenesis and screening. In earlier rounds, positions Q29 and N53 were mutated to positively charged Arg, a potential source of salt sensitivity. Therefore, the two sites were revisited with 19 AA substitution libraries and screened in a buffer approximating the intracellular salt composition of a mammalian cell (50 mM KH_2_PO_4_ pH 7.4) (*32*). Screening these libraries revealed that substituting His at residue 53 confers SNAP23 specificity to LC/A even in salt buffer, improving screening specificity 1300-fold compared to qmLC/A (**Fig. 2B-C. Fig. S4**). This selectant was termed the octuple mutant LC/A (omLC/A).

Prior work attributed substrate specificity in LC/A to primarily the alpha and beta binding exosites (*20*, *26*, *33*, *34*). These binding sites comprise the first and last interactions between substrate and protease, respectively. The results presented here identify a new series of residues between these exosites exerting control over SNAP23/SNAP25 specificity. We term these substrate control loops 20 (LC/A residues 26 to 29) and 50 (residues 52 to 56) (**Fig. 1B, D**).

Two categories of substitutions in these substrate control loops improved SNAP23 specificity. First, the enzyme’s overall charge became more positive. For instance, the loss of E55 and E56 in control loop 50 increased SNAP23 specificity up to two-fold (**Fig. 2C**). Similarly, substituting the positively charged Arg into sites across both control loops (A27, Q29, N53) improved SNAP23 specificity up to two-fold for each substitution. Increasing the positive charge in substrate control loop 20 likely accommodates D191 and D198 in SNAP23 (**Fig. S5**). Indeed, mutating SNAP25 residues to SNAP23 residues at these sites (E183D, T190D) reduces cleavage by wtLC/A up to 60% compared to cleavage of the native SNAP25 (*20*).

The second category of substitutions conferring greater SNAP23 specificity in these substrate control loops focuses on sidechain size. For example, substituting a small, hydrophilic serine for N26 in Round 3 increased specificity by nearly five-fold compared to the best Round 2 variant (**Fig. 2C**). Conversely, mutating the neighboring A27 to the larger Leu sidechain increased specificity 2.4-fold from Round 6 to Round 7. Position 26 could directly contact the SNAP23 substrate, whereas position 27 is directed towards control loop 50 (**Fig. S5**). Changing the size and flexibility of these residues therefore may reshape this SNAP-LC/A binding region to favor SNAP23 binding over SNAP25.

In addition to the discovery of substrate control loops 20 and 50 in LC/A, directed evolution revealed new information on established binding sites. At the alpha exosite, the SNAP25 residues in contact with LC/A are highly homologous to those in SNAP23, which suggests both substrates bind in a similar manner to this region. Indeed, generating libraries at alpha exosite residues in LC/A, such as K337, yielded few variants with proteolytic activity, and none with SNAP23 specificity (**Fig. S6**). Targeting residues near or within the beta exosite, however, was more productive. For example, libraries at N240 and S254 generated variants with up to three-fold improved SNAP23 specificity (**Fig. 2C**).

Comparing the enzyme kinetics for omLC/A and qmLC/A reveals the basis for its improved specificity. The main driver for omLC/A’s improved SNAP23 specificity is a 20-fold reduction in catalytic efficiency (*k*_cat_/*K*_M_) for cleaving SNAP25 (**Table 1, Fig. 3A-B**). Additionally, omLC/A improves the catalytic efficiency for cleaving SNAP23 by 6-fold. Notably, our directed evolution strategy assessed both SNAP23 and SNAP25 cleavage activities, driving these trends. The catalytic rate (*k*_cat_) for hydrolysis of SNAP25 is 40-fold lower for omLC/A compared to qmLC/A. In line with previous studies (*26*, *35*, *36*), the eight added mutations in omLC/A impact catalytic rate more than substrate affinity. Overall, the results represent a 120-fold improvement in SNAP23-specific catalytic efficiency from qmLC/A to omLC/A.

**Table 1.** Enzyme kinetics of LC/A variants. Kinetic constants of recombinantly expressed and purified LC/A variants were determined via the DARET assay (*23*) with modifications as described in the *Materials & Methods*. Values are means ± SEM (n=3) and und = undetectable.

|  | SNAP23 |  |  |  | SNAP25 |  |  |  |
| --- | --- | --- | --- | --- | --- | --- | --- | --- |
| | $k_{cat}$<br>(s<sup>-1</sup>) | $K_m$<br>(μM) | $V_{max}$<br>(mP*μM* s<sup>-1</sup>) | $k_{cat}/K_m$<br>(μM<sup>-1</sup>* s<sup>-1</sup>) | $k_{cat}$<br>(s<sup>-1</sup>) | $K_m$<br>(μM) | $V_{max}$<br>(mP*μM* s<sup>-1</sup>) | $k_{cat}/K_m$<br>(μM<sup>-1</sup>* s<sup>-1</sup>) |
| wtLC/A | und. | und. | und. | und. | 52 ± 2 | 3.4 ± 0.3 | 0.52 ± 0.02 | 15 ± 1 |
| qmLC/A | 3.3 ±<br>0.4 | 18 ± 4 | 0.033 ±<br>0.004 | 0.18 ±<br>0.05 | 140 ± 9 | 0.7 ± 0.2 | 1.40 ± 0.09 | 200 ± 60 |
| omLC/A | 8.1 ±<br>0.3 | 7.4 ±<br>0.6 | 0.081 ±<br>0.003 | 1.1 ± 0.1 | 3.2 ±<br>0.1 | 0.32 ±<br>0.09 | 0.032 ±<br>0.001 | 10 ± 3 |

**Fig. 3.**
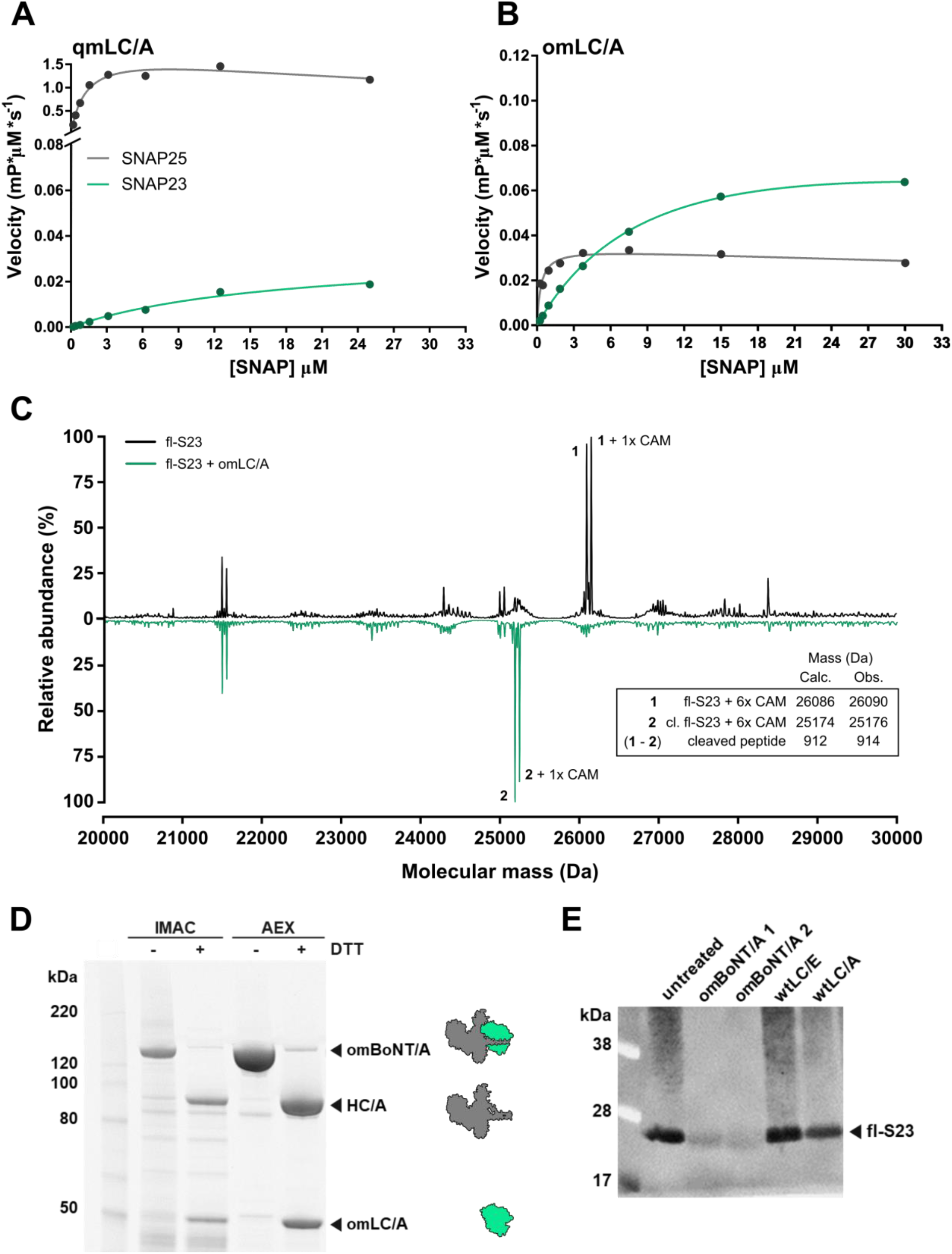
Characterization of omLC/A cleavage. (**A**) Rates of SNAP cleavage by batch-expressed, purified qmLC/A and (**B**) omLC/A at the indicated DARET substrate concentrations (n=3). (**C**) Deconvoluted ESI-MS of recombinant, full-length SNAP23 (fl-S23) treated with DTT and iodoacetamide to carbamidomethylate its cysteines (6x CAM). The mass spectrum of intact fl-S23 incubated with buffer (top, black) is compared to that of fl-S23 incubated with omLC/A (bottom, teal). Intact fl-S23 (**1**) and cleaved fl-S23 (cl. fl-S23, **2**) peaks are labeled. Additional marked peaks correspond to the masses of peaks **1** and **2** plus one additional CAM (+57 Da), likely resulting from overalkylation by iodoacetamide (*37*). (**D**) Recombinant omBoNT/A was purified by IMAC followed by anion exchange (AEX) chromatography. The omBoNT/A is ≈95% nicked upon DTT reduction as demonstrated by the presence of the HC/A and omLC/A bands. (**E**) *In vitro* cleavage of recombinant fl-S23 by two independent preparations of omBoNT/A (1 and 2) visualized with a C-terminal anti-SNAP23 antibody; before proteolysis, omBoNT/A was reduced with TCEP. The untreated, wtLC/A, and wtLC/E lanes provide negative controls.

Interestingly, the omLC/A variant exhibits strong dependence on Zn^2+^ for its substrate specificity, but not its activity. Without added Zn^2+^ during purification, the enzyme remains proteolytic, presumably due to endogenous Zn^2+^, but has a higher rate of cleavage for SNAP25 than SNAP23 (**Fig. S7**). The addition of Zn^2+^ (2 μM) during protein isolation reverses this specificity, and restores the enzyme’s preference for SNAP23. Importantly, once treated with Zn^2+^, the enzyme remains specific for SNAP23.

We next determined how omLC/A recognizes and cleaves the full-length SNAP23 (fl-S23) protein using mass spectrometry. Loss of the eight C-terminal residues of fl-S23 (RAKKLIDS) is observed upon treatment with omLC/A (**Fig. 3C**). This cleavage site in fl-S23 aligns with the expected SNAP25 site for wtLC/A hydrolysis (**Fig. 1D**). Therefore, the stringent cut-site specificity of this class of proteases is preserved in our engineered omLC/A variant for SNAP23; indeed, omLC/A fails to cleave the C-terminal domain of SNAP29 (**Fig. S8**), which is highly homologous to both SNAP23 and SNAP25 (**Fig. 1A**). The results suggest omLC/A interrogates its substrates and binds similarly to wtLC/A binding SNAP25 (as shown in **Fig. S5**).

We next incorporated omLC/A into the full-length BoNT/A holotoxin, termed omBoNT/A. Recombinant expression of omBoNT/A yielded a soluble 150 kDa polypeptide; as expected, reduction by DTT released the 100 kDa (HC/A) and the 50 kDa (omLC/A) proteins (**Fig. 3D**). Release of BoNT LC proteases through disulfide bond reduction is vital to successful cytosolic delivery and SNAP cleavage (*6*). The omBoNT/A retains its SNAP23 cleavage activity (**Fig. 3E**). Additionally, the omBoNT/A exhibits at least 25-fold reduced muscle paralysis associated with SNAP25 cleavage compared to wtBoNT/A (**Fig. S9**); the residual paralysis observed demonstrates successful cellular delivery of active omLC/A into motor nerve terminals. Furthermore, no systemic toxicity effects were observed for mice treated with 5 ng/kg omBoNT/A (**Fig S9**). Additional engineering of the omLC/A holotoxin is needed to enable efficient uptake into non-neuronal cells and to enable demonstration of SNAP23 cleavage activity *in vivo*.

This report demonstrates the plasticity of wtLC/A’s hyperspecific substrate preferences. Furthermore, omLC/A preserves wtLC/A’s vaunted selectivity for cleaving a single targeted peptide bond, with no additional cleavage products observed by MS. The eight mutations of omLC/A do not interfere with the omBoNT/A’s ability to bind and then release its peptide “belt,” as required for the holotoxin’s successful assembly and function. Thus, the successful incorporation of omLC/A into the BoNT holotoxin suggests that this reengineering will not interfere with its remarkable therapeutic properties of half-life and cell delivery.

The SNAP25 substrate wraps around nearly the entire circumference of LC/A, complicating rational attempts to alter substrate specificity. We address this with a novel fluorescence polarization-based platform that expands substrate screening from peptide-sized to include an entire domain of SNARE proteins. The approach offers general applicability, and can be applied to non-BoNT proteases. However, the SNARE proteins are well-suited to the extreme specificity testing demanded by the BoNT family proteases; for example, the SNAREs are structurally malleable. For non-BoNT proteases, which might not feature extremely long half-lives, catalytic rates should be considered with the substrate specificity index to identify the most active and selective proteases.

The results reported here can also guide future engineering of BoNTs. Our data suggests avoiding the alpha exosite for reengineering substrate specificity. However, pan-BoNT/A inhibitors (e.g., for anti-botulism therapies) could target the alpha exosite due to its recognition of a highly conserved region of SNAP substrates. Notably, substrate control loops 20 and 50 are structurally conserved amongst BoNT serotypes, yet feature divergent sequences (**Fig. S10**), which suggests their importance to directing the diverse substrate specificities of these proteases. Therefore, the strategy we describe could be broadly applicable.

BoNT represents a unique biotherapeutic as it catalytically ablates its target. Such capabilities are becoming increasingly important with PROTAC and other protease linkers emerging in pharmaceutical pipelines. We hope this work inspires new uses for BoNT proteases to treat a wide range of diseases.

## Acknowledgments

We thank Karen Brami-Cherrier for helpful conversations.

## Author contributions

G.A.W., A.D.B.-A., L.E.S., B.P.S.J. and S.M. conceptualized the project. R.P.D. and G.A.W. wrote the paper. R.P.D., G.S., M.H.F., L.Q.L., S.Z.M., E.J.V.-C., A.C.C., D.C.R., L.E.S. and G.A.W. designed experiments. R.P.D., H.M.I., G.S.S., M.H.F., L.Q.L., S.L., S.Z.M., E.J.V.-C., A.C.C., and T.T.M.N. performed the experiments. R.P.D., H.M.I., G.S.S., G.A.W., M.H.F., L.Q.L, S.L., S.Z.M., E.J.V.-C., and A.C.C. analyzed the results. A.D.B.-A, L.E.S., E.J.V.-C., and S.Z.M. edited the manuscript.

## Data and materials availability

All data needed to evaluate the conclusions in the paper are present in the paper and/or the Supplementary Materials. Additional data related to this paper may be requested from the authors.

## Supplementary Materials

Materials and Methods

Figs. S1 to S10

References (*38-41*)

## Supplementary Information

## Materials and Methods

### LC/A variants

#### Construction of the qmLC/A encoding plasmid

The gene encoding the qmLC/A variant (E148Y, K166F, S254A, G305D) was constructed through overlap extension PCR using the gene encoding wild-type LC/A (WT, amino acids 1 to 430) as the template. The resulting PCR products were *DpnI*-digested, column purified and concentrated, then ligated and subcloned via Gibson Assembly into a pET-29b(+) vector featuring a C-terminal His_6_ tag for recombinant protein expression and purification. Sanger sequencing (Genewiz LLC) confirmed construction of qmLC/A-pET-29b(+).

#### Batch expression and purification of LC/A variants

The LC/A-pET-29b(+) WT and mutant plasmids were transformed into BL21 (DE3) *E. coli* (One Shot™ Star™, ThermoFisher) cells and spread on LB_kan_ agar plates at 37 °C overnight. A single colony was used to inoculate an LB_kan_ seed culture incubated for 6 to 18 h at 37 °C with shaking at 225 rpm. The LB_kan_ expression cultures were supplemented with glucose (1% w/v) and inoculated with seed culture (1:1000), then incubated at 37 °C, with shaking at 225 rpm until the culture reached an OD_600_ of 0.6. The LC/A protein production was induced by addition of isopropyl β-D-1-thiogalactopyranoside (IPTG, 1 mM final concentration) and incubated at 25 °C with shaking at 225 rpm for 20 to 24 h. The cells were harvested by centrifugation (6084 × g, 4 °C, 20 min) and resuspended while on ice in LC Lysis Buffer (100 mM HEPES, 25 mM imidazole, 500 mM NaCl, pH 7.4) for storage at −80 °C or immediately subjected to lysis. The soluble protein was extracted from cells via sonication followed by centrifugation (26,891 × g, 4 °C, 45 min), and the supernatant then batch bound overnight at 4 °C to nickel-charged IMAC resin (Profinity, BioRad) which had been equilibrated with LC Lysis Buffer. The LC/A variants were purified via a gravity column, which was first washed with 20 column volumes (c.v.) of LC Lysis Buffer, then 12 c.v. LC Lysis Buffer supplemented with 50 mM imidazole, and finally eluted with 16 c.v. LC Lysis Buffer supplemented with 500 mM imidazole, collecting four 4-mL fractions. The fractions were analyzed by SDS PAGE (12% acrylamide) and visualized by Coomassie blue stain. LC/A-containing fractions with a purity of 95% or higher were combined and dialyzed into chilled Storage Buffer at 4 °C. After sterile filtration through a 0.22 μm filter, the purified LC/A variants were subjected to BCA assay to quantify protein concentration through comparison to BSA standards (Pierce™ BCA Assay, ThermoFisher). The proteins were stored at 4 °C before assays.

### Fluorescence polarization substrates

#### Expression and purification of SNAP23 & SNAP25 substrates

Substrates for the depolarization after resonance energy transfer (DARET) assay comprising of the C-terminal domain of SNAP23 (amino acids 137 to 211) or SNAP25 (amino acids 134 to 206) fused between GFP and BFP were subcloned into pET-28c(+) via ligation independent cloning. The DARET-pET-28 plasmids were transformed into BL21 (DE3) *E. coli* cells and spread on LB agar plates supplemented with 40 μg/mL kanamycin (LB_kan_) then incubated at 37 °C overnight. A single colony was used to inoculate an LB_kan_ seed culture and incubated for 6 to 18 h at 37 °C with shaking at 225 rpm. Next, LB_kan_ expression cultures were inoculated with 1:1000 of the seed culture and incubated at 37 °C with shaking at 225 rpm until the cultures reached an OD_600_ of 0.6. The DARET substrate production was induced through addition of IPTG (final concentration of 0.5 mM) and incubated at 16, 18, or 22 °C with shaking at 225 rpm for 18 to 24 h. The cells were harvested by centrifugation (6084 × g, 4 °C, 20 min) and resuspended while on ice in DARET Lysis Buffer (25 mM HEPES, 300 mM NaCl, pH 8.0) for storage at −80 °C or immediately subjected to lysis. The soluble protein was extracted from cells via sonication followed by centrifugation (26,891 × g, 4 °C, 45 min), and the supernatant was mixed with urea (0.5 M final concentration) before batch binding overnight at room temperature (RT) to nickel-charged IMAC resin (Profinity, BioRad) that had been previously equilibrated with the DARET Lysis Buffer. The DARET substrates were purified in a gravity column, first washing with 20 (c.v.) DARET Lysis Buffer, then 12 c.v. of DARET Lysis Buffer supplemented with 20 mM imidazole, and finally eluted with 16 c.v. of DARET Lysis Buffer supplemented with 250 mM imidazole. During elution, fractions with visibly green DARET substrates were collected. These fractions were combined and transferred to dialysis tubing (SnakeSkin, ThermoFisher) and dialyzed into chilled Storage Buffer (50 mM HEPES pH 7.4) at 4 °C. After sterile filtration through a 0.22 μm filter, the purified DARET substrates were subjected to BCA assay to quantify protein concentration through comparison to BSA standards. The DARET substrates were then aliquoted, protected from light, and stored at 4 °C or flash frozen for storage at −80 °C before assays.

#### Construction, expression, and purification of SNAP29 substrate

SNAP29 DARET was constructed by inserting a gene fragment of human SNAP29 (amino acids 185 to 258, Genewiz) between eGFP and eBFP2 via overlap extension PCR, then cloning into pET-28c(+) through Gibson assembly. The SNAP29 DARET-pET28-c(+) construct was confirmed by Sanger sequencing (Genewiz). The SNAP29 DARET-pET-28 plasmid was transformed into BL21 (DE3) *E. coli* cells and spread on an LB_kan_ agar plate then incubated at 37 °C overnight. The SNAP29 DARET substrate was expressed as described for the SNAP23 and SNAP25 DARET substrates with minor alterations (OD_600_ of 0.8, 0.25 mM final IPTG). Purification and storage of the SNAP29 substrate were performed as described for SNAP23 and SNAP25 DARET substrates.

### High-throughput specificity screens

#### Library generation

The qmLC/A encoding plasmid served as a template for the first round of mutagenesis, and the qmLC/A provided a positive control and standard for each screen of SNAP23/25 specificity. The PCR products were *DpnI*-digested, column purified and concentrated, and then subcloned into pET-29b(+) via Gibson Assembly and evaluated by Sanger sequencing. The error-prone PCR (epPCR) library of the gene encoding qmLC/A was created with the GeneMorph II kit (Agilent) according to the manufacturer’s instructions. The Round 1 epPCR library yielded the following mutants having SNAP23/SNAP25 specificity greater than qmLC/A: E201D/D203V, N240S, K364R/Y387N. Site-directed mutagenesis libraries were generated with overlap extension PCR using oligonucleotides featuring degenerate sequences at selected residues. In Round 2, the saturated amino acid substitution library (20 amino acids) was constructed using Tang’s mutagenesis method (*39*) at position 240. In Rounds 3 to 6 and part of Round 7, iterative saturation mutagenesis using an NDT codon (encoding Cys, Asp, Phe, Gly, His, Ile, Leu, Asn, Arg, Ser, Val, and Tyr) was employed at residues N26, A27, G28, Q29, M30, T52, N53, E55, E56, E171, S143, Q162, M253, T327, and K337. These rounds yielded the N26S, Q29R, N53R, and E55V variants. In the second part of Round 7 and the entirety of Round 8, libraries featuring NDT and VHG codons (substitution with all AAs except Trp) were constructed at residues Q29, N53, Y148, F166, A254, and D305. These rounds yielded the N53H, A254L, and D305G variants. Each saturation library was subject to Q_pool_ analysis for quality control (*30*); libraries with a Q_pool_ <0.7 were subcloned again from PCR fragments and further analyzed before screening.

#### Screening for SNAP23/SNAP25 specificity

The qmLC/A libraries were transformed into BL21 (DE3) *E. coli* cells then spread onto LB_kan_ agar plates for overnight incubation at 37 °C. Single colonies were selected and inoculated single 96-well deep-well plate (DWP) seed cultures (300 μL LB_kan_ per well), then incubated at 37 °C with shaking at 225 rpm for 18 h. A qmLC/A and uninoculated culture were included as single wells on each DWP as positive and negative controls, respectively. An expression culture (630 μL LB_kan_ per well) was subsequently inoculated with 20 μL of the seed culture and incubated for 3 h at 37 °C with shaking at 225 rpm. To generate glycerol stocks, the cells remaining from the seed culture DWP were harvested via centrifugation (2056 × g, 4 °C, 1 h), then resuspended in 50 to 100 μL ultrapure glycerol (50% in autoclaved nanopure water) and transferred to a 96-well plate for storage at −80 °C. The expression DWP culture was chilled on ice for 10 min, then induced by adding IPTG (1 mM final concentration) before incubating for 22 to 24 h at 23 °C with shaking at 900 rpm. The cells were harvested via centrifugation (2056 × g, 4 °C, 1 h) then stored at −80 °C for 20 min or overnight. For screening, cells were thawed at RT, then lysed at 23 °C with shaking at 500 rpm for 30 min in DWP Lysis Buffer: 100 μL per well of SoluLyse (Genlantis) plus benzonase nuclease (75 U/mL, NEB). The insoluble debris was removed via centrifugation (2056 × g, 4 °C, 1 h), then an aliquot (75 μL) of each lysate was transferred to a 96-well plate (Celltreat). Lysates were diluted 1:100, 1:200, 1:400, or 1:600 in Assay Buffer (50 mM HEPES, 0.05% v/v Tween, pH 7.4) or Intracellular Buffer (50 mM KH_2_PO_4_, pH to 7.4 with KOH) before screening. Recombinantly expressed and purified SNAP25 DARET substrate was diluted to 3 μM in Assay Buffer or Intracellular Buffer, then 100 μL substrate added to each well of a 96-well flat, black, non-binding surface microtiter plate (Corning). From the diluted lysate plate, 50 μL of the blank was added to its corresponding well in the black plate and used to optimize the gain and Z position of a Spark fluorescence polarization plate reader (Tecan). The sample was excited with polarized light at 380 nm with a 20 nm bandwidth, written here as 380(20) nm, and the polarized emission detected at 535(25) nm. For the remaining 95 wells, 50 μL from each well of the lysate dilution plate was added to the black plate containing substrate and the entire plate screened kinetically for 50 min to 14 h at 28 ± 1 °C. The assay steps were then repeated for 3 μM SNAP23 DARET using the same lysate dilution plate. The final concentration of each substrate screened was 2 μM. The changes in polarization over time were visualized using Prism (GraphPad) and initial rates (V_0_) were derived from fitting trendlines to the initial, linear portion of the raw data. The rates of negative controls (no enzyme) for each DARET substrate were subtracted from the rates of each variant to account for nonenzymatic changes in polarization. The specificity indices were calculated via the ratio of SNAP23 and SNAP25 initial rates for each clone according to Eq. 1.

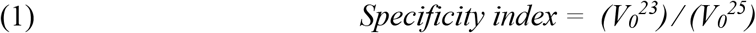

In Rounds 2 to 8, LC/A variants with specificity indices at least 1.5 times higher than qmLC/A or the most specific variant from the previous round of directed evolution were subject to further screening in triplicate. From the glycerol stocks, the variants and controls (qmLC/A, last best variant) were streaked onto LB_kan_ plates for overnight incubation at 37 °C. Three colonies from these streaks were used to inoculate three wells of a seed culture DWP, then three wells of an expression culture DWP for LC/A variant production, harvesting, lysis, and screening as described above. The specificity index of each well was calculated first, then indices averaged together for the same variant. LC/A variants demonstrating consistent, improved specificity over the last best variant (>1σ, where σ = standard deviation) were selected as starting points for the next round of mutagenesis and screening.

### Biochemical characterization

#### Mass spectrometry

To determine the SNAP23 cleavage site of the LC/A variants, recombinant full-length SNAP23 (fl-S23) with N-terminal hexa-His and TEVp tags was expressed and purified to >95% purity by SDS PAGE, then dialyzed into 20 mM Tris pH, 8.0 at 4 °C overnight. For mass spectrometry (MS) assays, fl-S23 was treated with dithiothreitol (7.5 mg/mL) at 80 °C for 10 min followed by iodoacetamide (9 mg/mL) at 25 °C for 1 h in the dark. This treatment functionalized the six free cysteines between the N- and C-terminal domains to prevent inter- and intra-disulfide bonding, thus simplifying any subsequent mass spectra. The functionalized fl-S23 was then incubated overnight at RT with the omLC/A variant at final concentrations of 2 μM substrate, 50 nM protease in Storage Buffer. The negative control was fl-S23 incubated with Storage Buffer. The samples were analyzed by triple-quad liquid chromatography/electrospray ionization in a Xevo TQ-S mass spectrometer (Waters), and the spectra were deconvoluted in MassLynx using the MaxEnt1 algorithm.

#### Kinetics

LC/A variant kinetic parameters (*K*_m_, *k*_cat_) were measured by monitoring the increase in millipolarization units (mP) in a Spark M10 fluorescence polarization plate reader (Tecan) at 28 ± 1 °C over 2 h, excited with polarized light at 380(20) nm and detected with polarized emission at 535(25) nm. The Michaelis-Menten kinetic experiments for DARET SNAP23 and SNAP25 substrates employed a substrate concentration range of 0.2 to 30 μM, with a fixed concentration of 10 nM LC/A variant in 50 mM KH_2_PO_4_ 0.2 nM ZnCl_2_ pH 7.4. Each concentration of substrate with LC/A variant or buffer (blank or negative control) was assessed in triplicate (n=3). The average change in polarization over time observed for the buffer was subtracted from the rate observed for each LC/A-incubated well at the substrate concentration; the change in polarization monitored by the blank was exceptionally low. The initial rates (V_0_) were derived by fitting the first 25 to 35 min of blank-subtracted rates to a linear regression model in Prism (GraphPad) and calculating the slope. Next, V_0_ values were multiplied by their corresponding substrate concentrations as previously described (*23*), and fit to Michaelis-Menten curves in Prism. Kinetic parameters for wild-type LC/A were derived from a *k*_cat_ model, and parameters for qmLC/A and omLC/A were fit to a substrate inhibition model. Goodness of fit (R^2^) values for wtLC/A (0.99), qmLC/A (0.97 for SNAP25, 0.99 for SNAP23), and omLC/A (0.87 for SNAP25, 0.99 for SNAP23) data guided which models were used. The *k*_cat_ values were derived by dividing V_max_ by the concentration of protease. For all assays, the blank with the highest substrate concentration was used to optimize the gain and Z-position of the plate reader before addition of protease.

#### Molecular modeling/visualization

Molecular graphics and structural analyses were performed with UCSF Chimera.

### Full-length omBoNT/A holotoxin

#### Batch expression and purification of omBoNT/A

The omBoNT/A variant was expressed in 1.5 L of Terrific Broth supplemented with 1% glucose (w/v) until the culture reached an O.D. of 0.6. The culture was then induced with 0.2 mM IPTG and incubated at 16 °C, 265 rpm for 20 hours. After cell harvest and lysis, omBoNT/A was purified by IMAC on MagneHis™ resin followed by anion exchange chromatography on a Hitrap Q HP column. Pooled anion exchanged fractions were exchanged into 50 mM Tris pH 8.0, 120 mM NaCl, 0.1 mM ZnCl2 and 5% PEG (v/v) 400 (#8074850050, Sigma). Fractions were analyzed for nicking in the presence or absence of 100 mM DTT by SDS PAGE, staining with Spyro Ruby.

#### SNAP23 *in vitro* cleavage assay

Human SNAP23 *in vitro* cleavage was evaluated by incubating 30 μg of human recombinant SNAP23 protein (Novus Biologicals, NBC1-18347) with 400 nM of either wild type LC/A, wild type LC/E, or reduced full length omBoNT/A at 37°C for 1 hour in PBS, pH 7.2 (ThermoFisher, 20012-043). The omBoNT/A holotoxin was reduced by incubation with 2 mM TCEP (Tris-(2-carboxyethly) phosphine, hydrochloride (ThermoFisher, T2556) at 37°C for 4 hours. The reaction was quenched by addition of NuPAGE™ LDS Sample Buffer (ThermoFisher, NP0008) and the amount of SNAP23 was assessed by Anti-SNAP-23 Western blot analysis. Reaction samples (~24 μg) were separated by SDS-PAGE (12% acrylamide) and transferred to nitrocellulose membrane, 0.45 μm pore size (ThermoFisher, LC2001) in 1X Transfer buffer (ThermoFisher, NP006) containing 20% v/v methanol (Honeywell, 230-4). Membranes were blocked in 2% ECL Prime™ Blocking Agent (GE Healthcare, RPN418) in TBST (Tris-Buffered Saline (Bio-Rad, 170-6435) with 0.1% Tween-20 (Bio-Rad, 161-0781)) for 1 h at RT. Intact SNAP-23 protein was detected with anti-SNAP23 polyclonal antibody (Synaptic Systems, 111203) diluted 1:1000 in 2% blocking buffer. Blots were incubated overnight with primary antibody at 4°C with gentle agitation. Blots were washed in TBST and the bound antibody was detected after 1 h incubation at RT with HRP-Goat Anti-Rabbit IgG (H-L) (ThermoFisher, G21234) diluted 1:4000 in 2% blocking buffer. After final washes in TBST, the membranes were reacted with Pierce™ ECL Plus Western Blotting Substrate (ThermoFisher, 32134) and scanned using the Typhoon 9410 Imager (GE Healthcare).

#### Mouse digital abduction (DAS) assay

All procedures were approved by AACUC (approved protocol #225-100051-2019). The DAS assay was performed as previously reported (*42, 43*). In summary, female CD-1 mice (Charles River), with an average weight of 30.2 g and age range of 6−10 weeks old, were used in this study. The omBoNT/A and purified BoNT/A1 neurotoxin (Metabiologics Inc., referred to as wtBoNT/A) were diluted in 0.5% human serum albumin in 0.9% saline (Fresenius Kabi, 918620). For the assay, 0.005 mL of each diluted holotoxin were injected in the right gastrocnemius muscle. Three mice per dose (n=3) were tested in triplicates (N=3). The DAS, well-being score, and weight were recorded daily for 4 days. The results were plotted using Prism (GraphPad). Each mouse’s well-being was scored on a 4-point system (0 = activity level normal; 1 = slightly diminished activity level and/or slight weight loss (5 - 10%); 2 = moderately diminished activity level and moderate weight loss (10 - 15%); 3 = severely diminished activity level, little to no reaction to outside stimuli, inability to ambulate, agonal or labored respiration).

## Supplemental Figures

**Figure S1.**
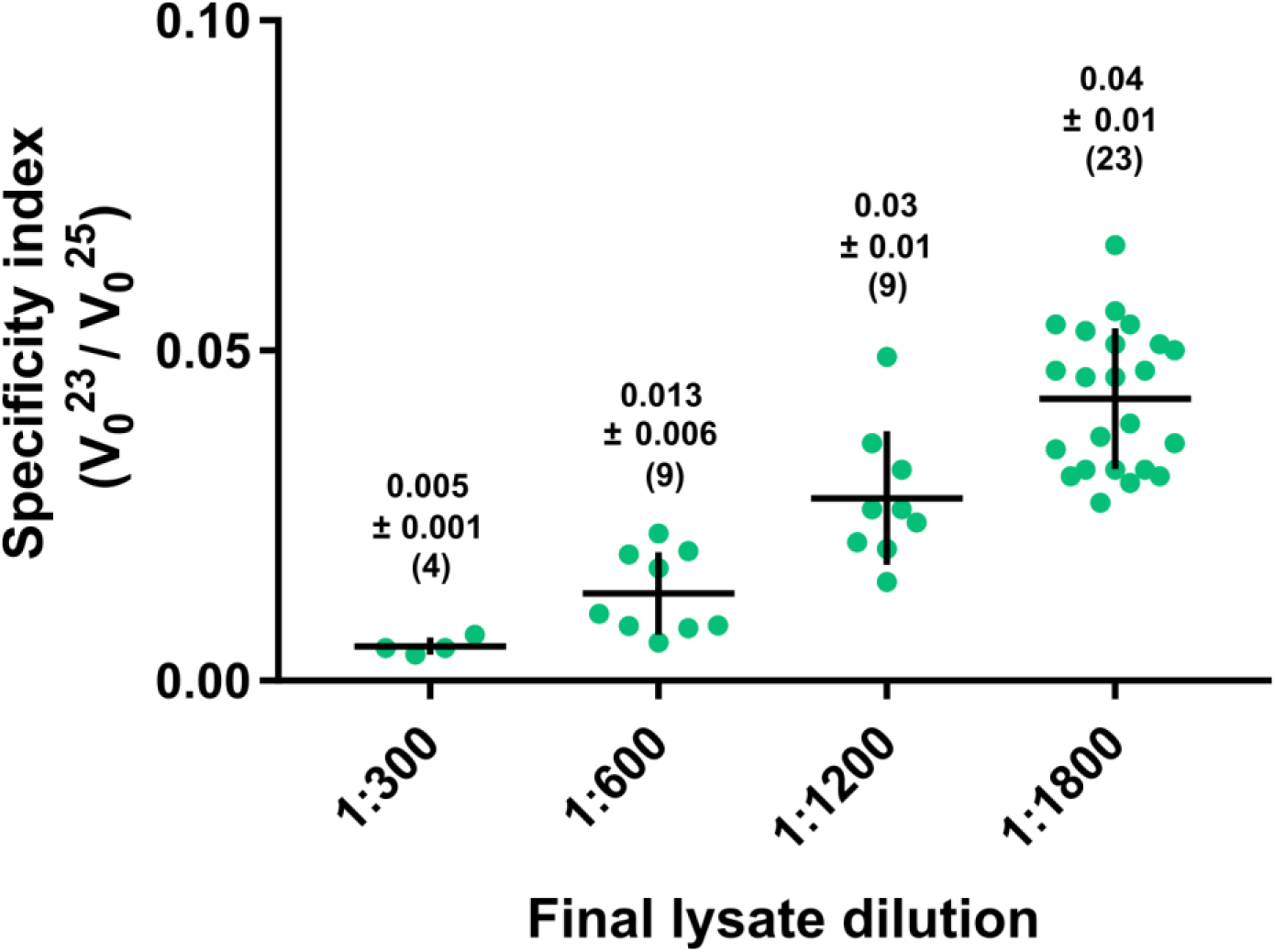
The substrate specificity of the qmLC/A variant. As shown by the specificity index (V_0_^23^/V_0_^25^) at 2 μM substrate, qmLC/A in cell lysate is highly specific for SNAP25. However, diluting the cell lysate increases the SNAP23 specificity of qmLC/A. Each point represents a different technical or biological replicate at the indicated final lysate dilution. The average value and standard deviation are shown as numbers above each dilution with the total number of data points in parentheses. Horizontal lines indicate average value and vertical lines indicate one standard deviation. The four dilutions yielded significantly different specificity indices as analyzed by an unpaired t-test in GraphPad Prism (*p* = 0.001).

**Figure S2.**
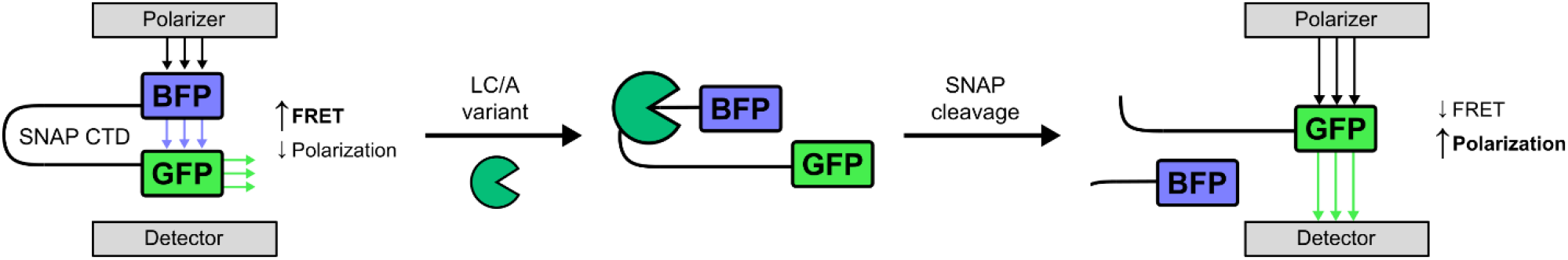
Depolarization After Resonance Energy Transfer (DARET) assay. Cartoon representing the DARET assay for SNAP cleavage by LC/A variants. CTD, C-terminal domain, corresponds to residues 137 to 211 of SNAP25 or residues 134 to 206 of SNAP23.

**Figure S3.**
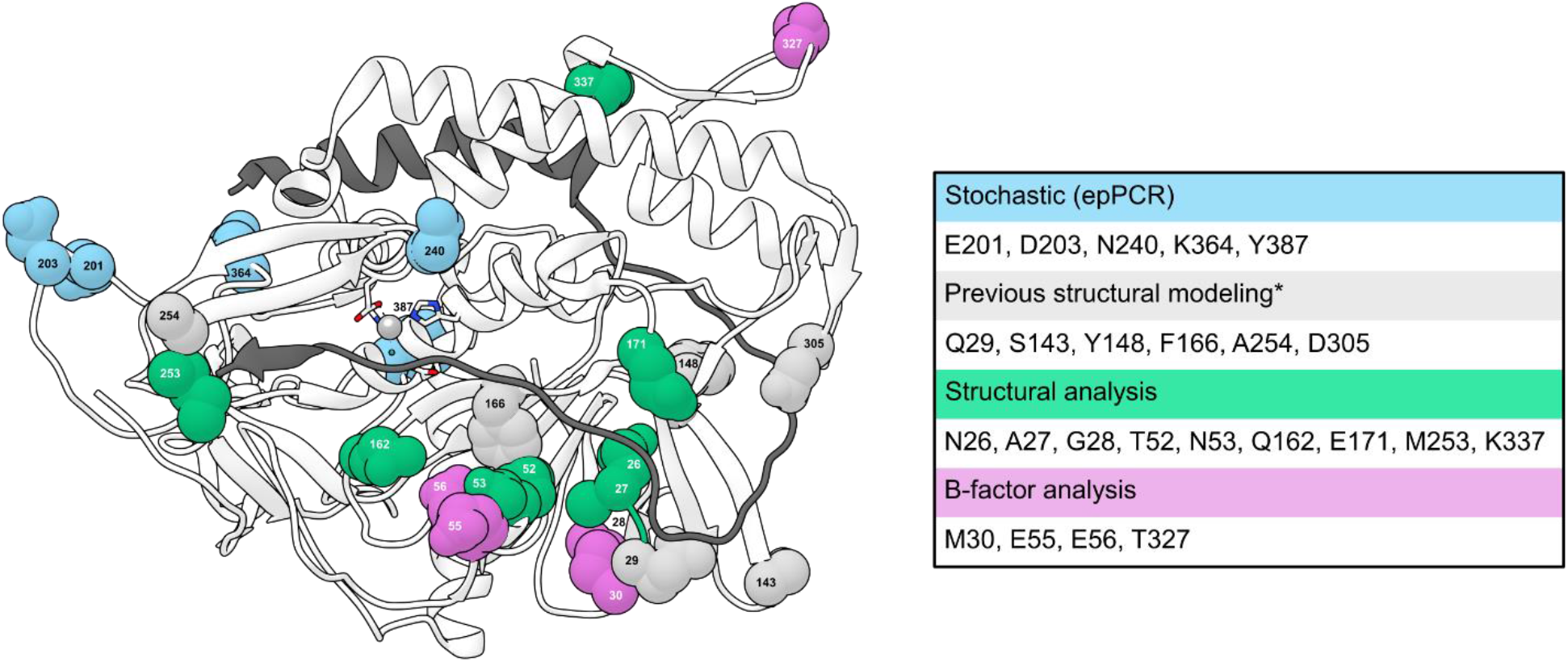
Residues targeted and techniques used for directed evolution. Residues highlighted in blue were identified via stochastic methods (error-prone PCR, epPCR). Gray residues were mutated with the NDT and VHG codons (all AAs substituted except Trp). Sites directing specificity in teal were discovered through structural analysis of the SNAP25-LC/A complex (1XTG, shown here), and were mutated with the NDT codon (12 AAs substituted: Cys, Asp, Phe, Gly, His, Ile, Leu, Asn, Arg, Ser, Val, Tyr). Purple residues were chosen based upon B-factor analysis, and were mutated with the NDT codon. Position 240 was substituted with all 20 AAs using Tang’s mutagenesis method. Asterisk (*) indicates the residues reported by Binz et al. (*21*).

**Figure S4.**
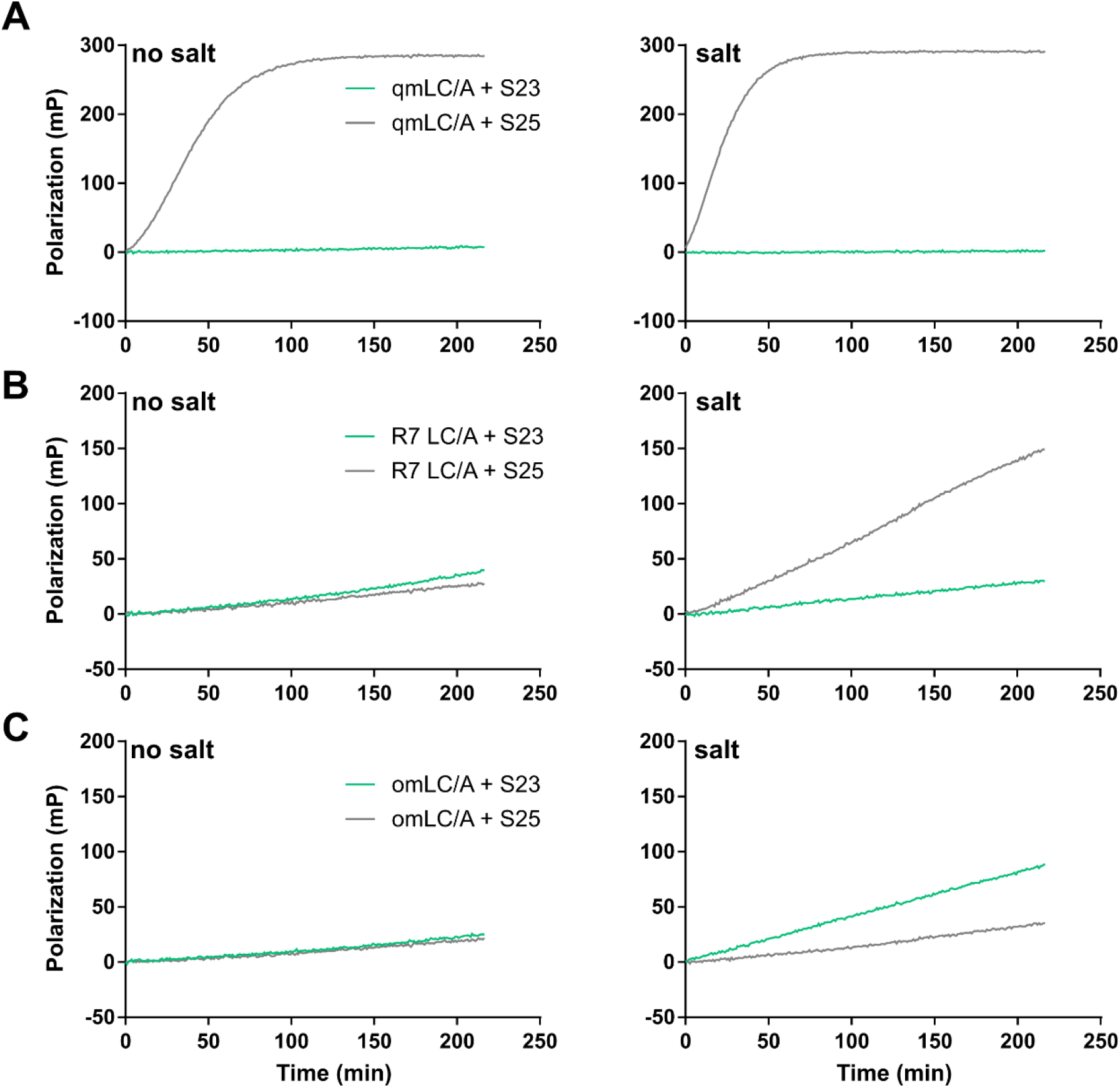
Salt sensitivity of LC/A selectants. In Round 8 of directed evolution, diluted lysates (1:1800) containing qmLC/A, omLC/A, or the most SNAP23 specific variant from Round 7 (R7) were tested for SNAP cleavage in “salt-free” (50 mM HEPES pH 7.4, 0.05% Tween) or salt (50 mM KH_2_PO_4_ pH 7.4) buffered conditions. (**A**) The qmLC/A variant is highly specific for SNAP25 in both no salt and salt conditions. (**B**) The R7 LC/A variant is specific for SNAP23 in no salt conditions but becomes specific for SNAP25 in salt. (**C**) The omLC/A variant is specific for SNAP23 in no salt and salt conditions. Each line depicts the mean of three technical replicates after subtracting a no-enzyme blank.

**Figure S5.**
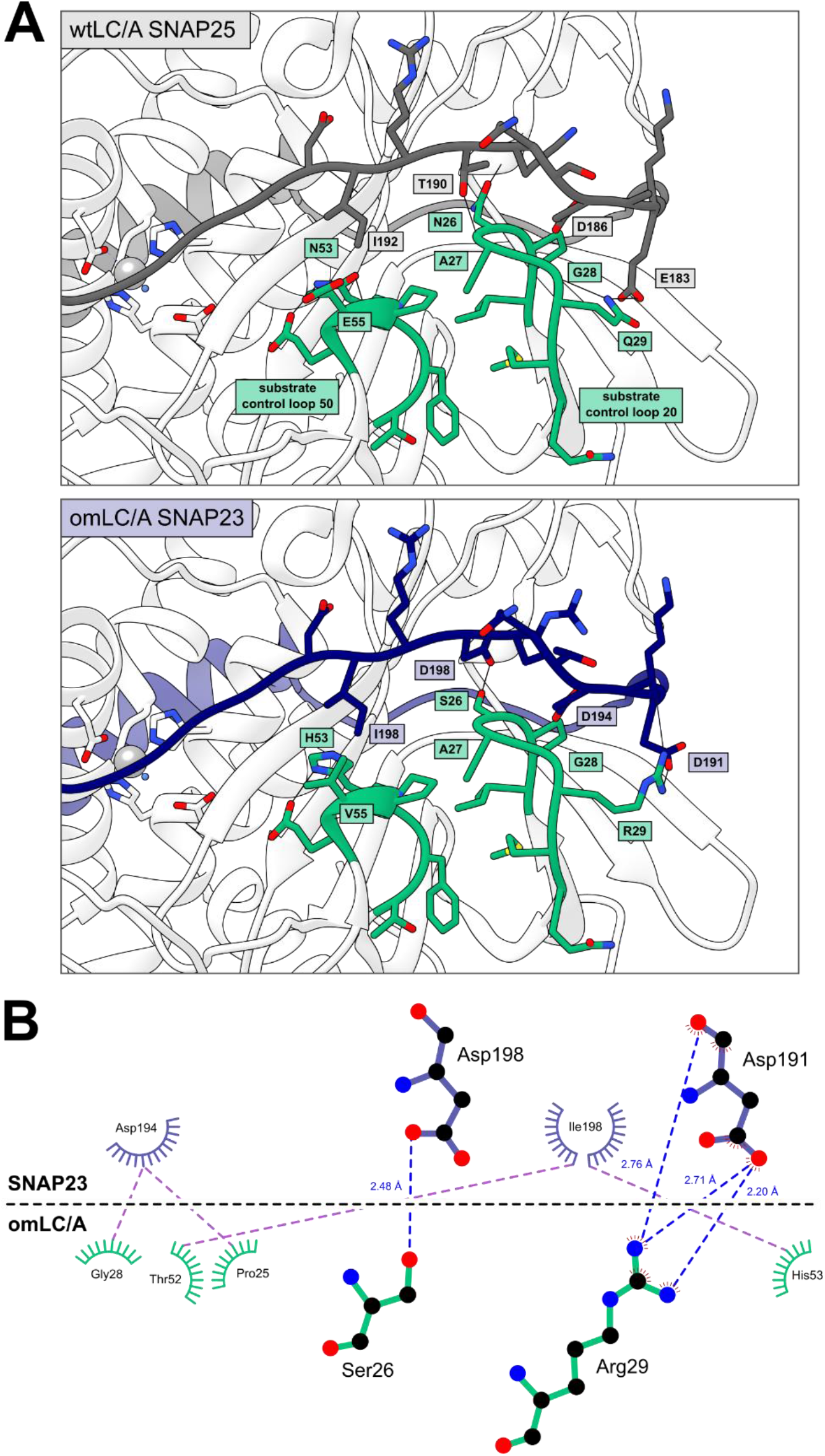
Substrate control loops in LC/A. (**A**) Residues occupying substrate control loops 20 and 50 are highlighted for wtLC/A (top) and omLC/A (bottom, computational modeling). (**B**) Putative modeled protein-protein interactions between SNAP23 sidechains (navy) and omLC/A substrate control loop residues (teal) (LigPlot^+^ software). Hydrophobic interactions are depicted in purple, and putative polar contacts are shown in blue.

**Figure S6.**
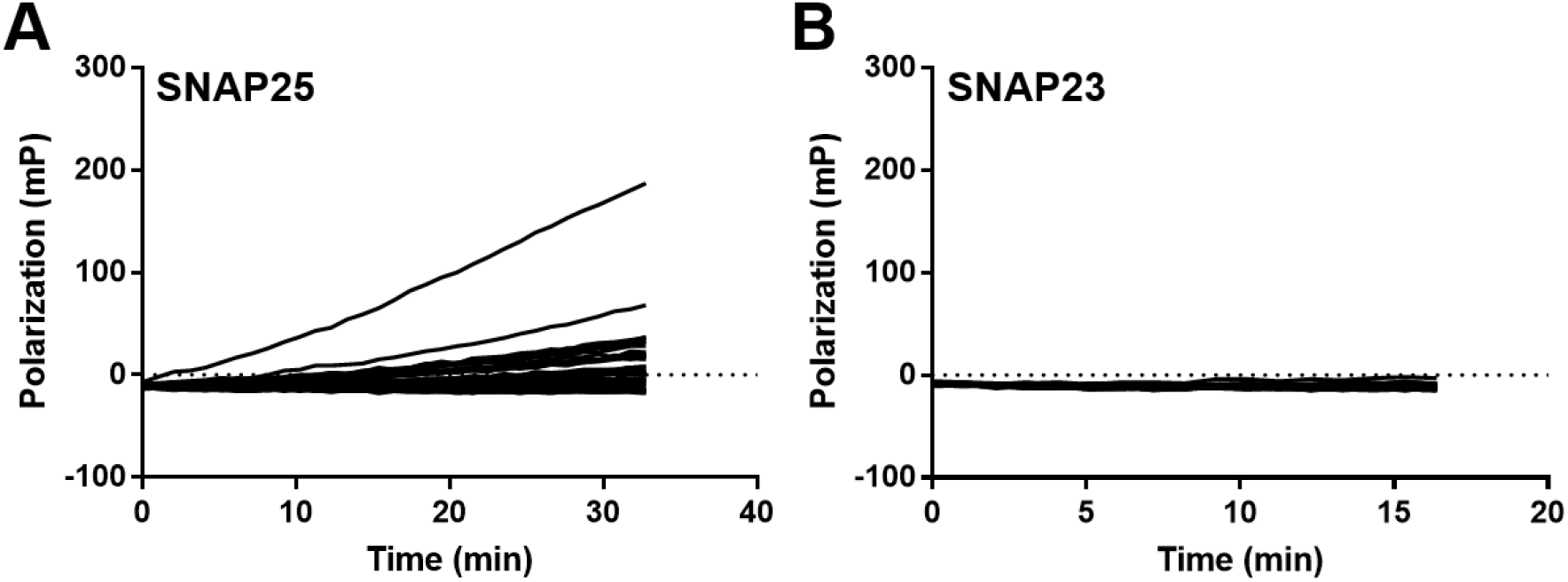
SNAP cleavage activity of K337NDT clones. In Round 6 of directed evolution, 34 variants from the alpha exosite K337NDT LC/A library were tested for (**A**) SNAP25 or (**B**) SNAP23 cleavage in the DARET assay. Each line represents the diluted lysate (1:1800) of a single well of a 96 deep-well plate.

**Figure S7.**
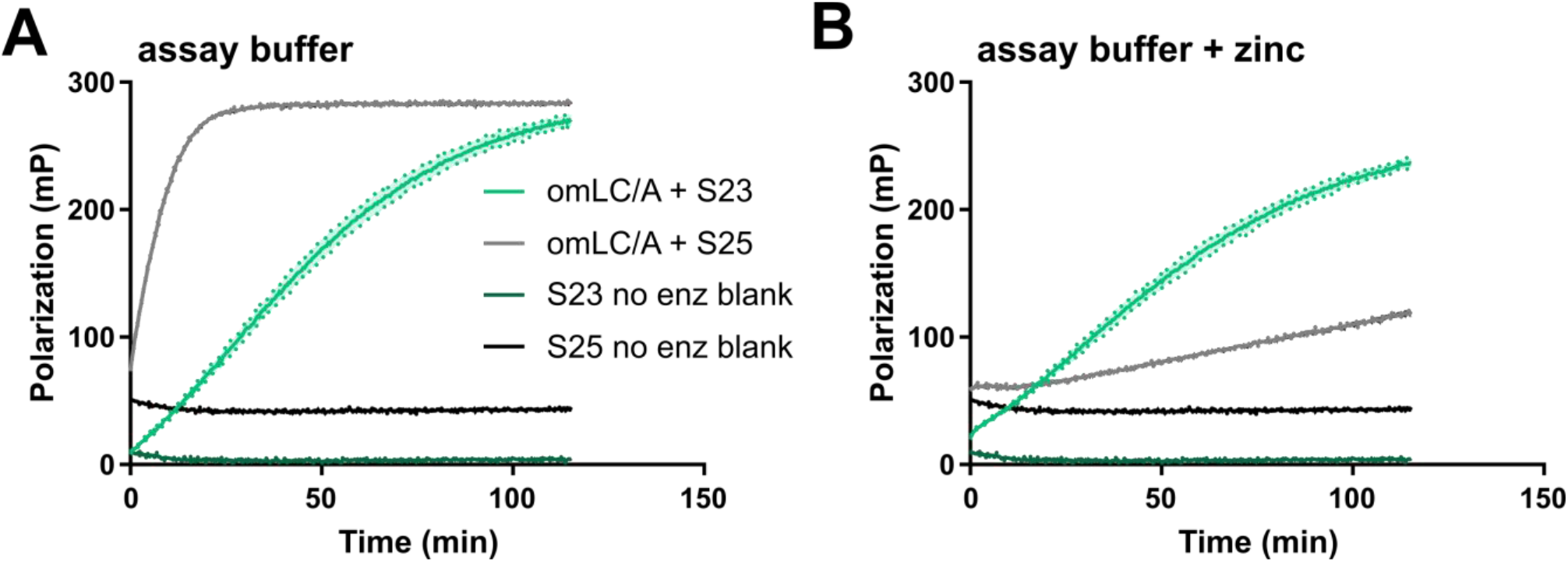
Zinc dependence of omLC/A substrate specificity. (**A**) Batch-expressed, IMAC-purified fractions of omLC/A were dialyzed into either Assay Buffer (50 mM HEPES pH 7.4) or (**B**) Assay Buffer plus zinc (50 mM HEPES, 2 μM ZnCl_2_ pH 7.4) for 20 h at 4 °C before characterizing SNAP cleavage by the DARET assay (no enz blank = no enzyme blank negative control). Typically, zinc is not supplemented during the purification of wtLC/A. Each solid line depicts the mean of three technical replicates (50 nM enzyme, 2 μM substrate). The dotted lines and the shaded area depict the standard deviation around the mean.

**Figure S8.**
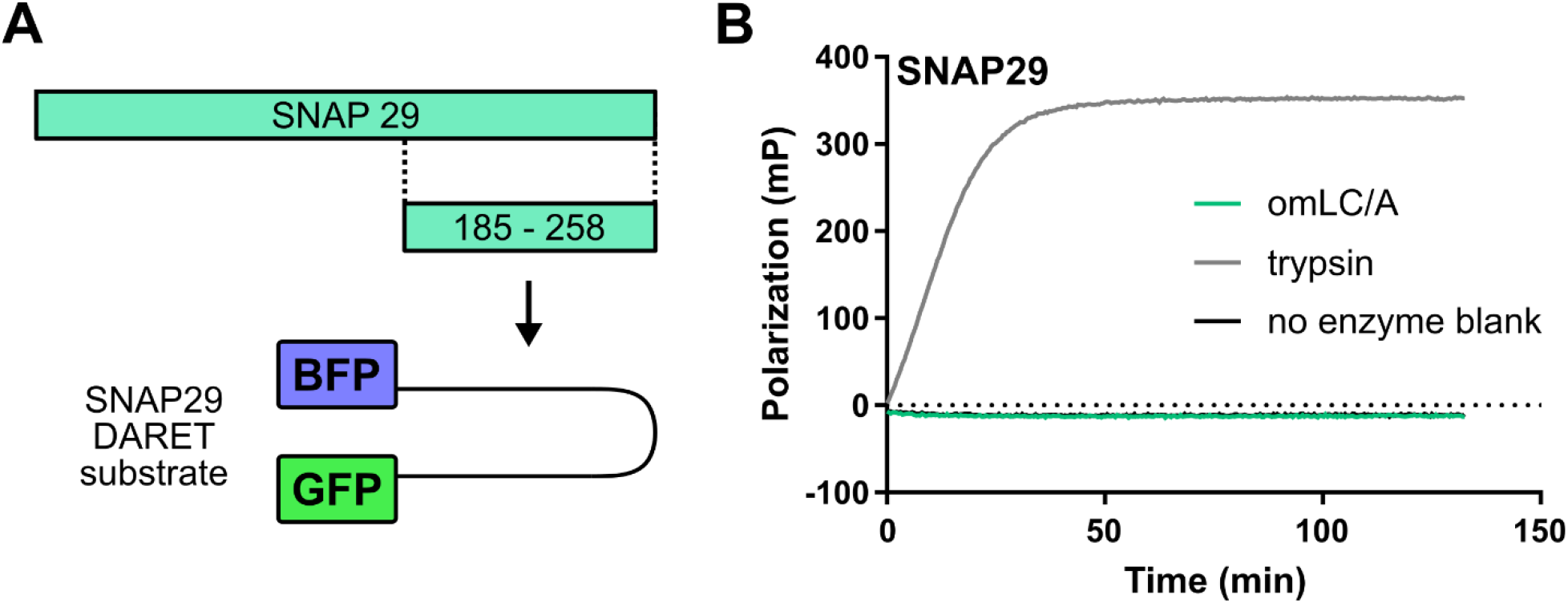
Cleavage of related SNARE SNAP29 by omLC/A. (**A**) Design of SNAP29 DARET substrate for fluorescence polarization assays of proteolytic cleavage, featuring the 73 C-terminal residues of SNAP29. (**B**) The omLC/A variant exhibits no SNAP29 cleavage in the DARET assay. SNAP29 is the closest related SNAP family member to SNAP25 and SNAP23. Trypsin provides a positive control for SNAP29 cleavage. Each solid line depicts the mean of three technical replicates (50 nM enzyme, 300 nM substrate). The dotted lines and the shaded area depict the standard deviation around the mean; however, the standard deviation for each condition tested is too small to be visible on the graph.

**Figure S9.**
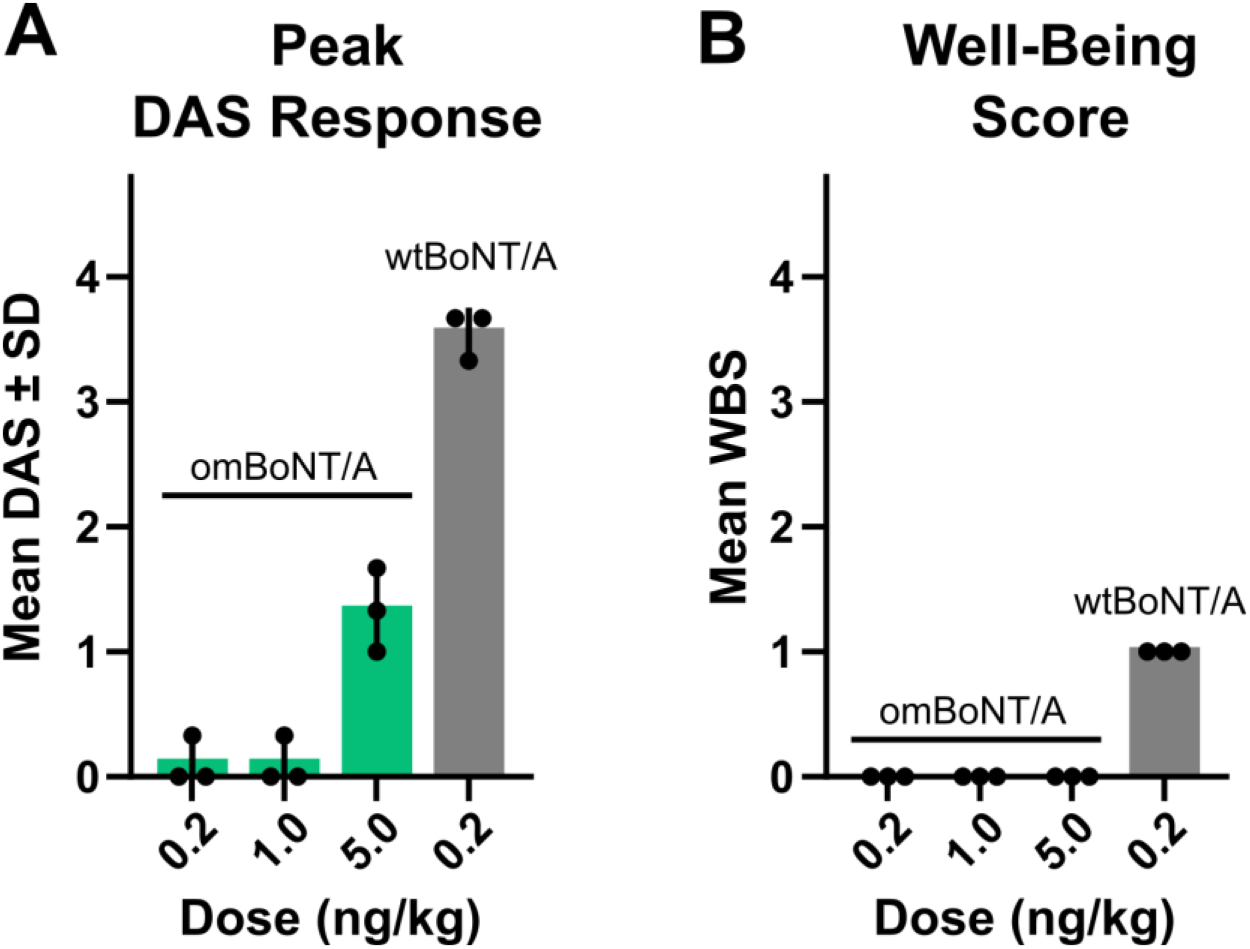
In vivo evaluation of neuromuscular paralysis of omBoNT/A compared to wtBoNT/A. (**A**) Neuromuscular paralysis measured by peak DAS effect in mice (n=3/dose; N=3) injected with omBoNT/A (0.2, 1, or 5 ng/kg) or wtBoNT/A (0.2 ng/kg). The paralytic response is mediated by cleavage of SNAP25. (**B**) Mice injected with omBoNT/A (n=3/dose; N=3) had no changes in the well-being score (WBS) over four days. These results confirm reduced SNAP25 activity *in vivo* and demonstrate that omBoNT/A does not result in increased overt toxicity.

**Figure S10.**
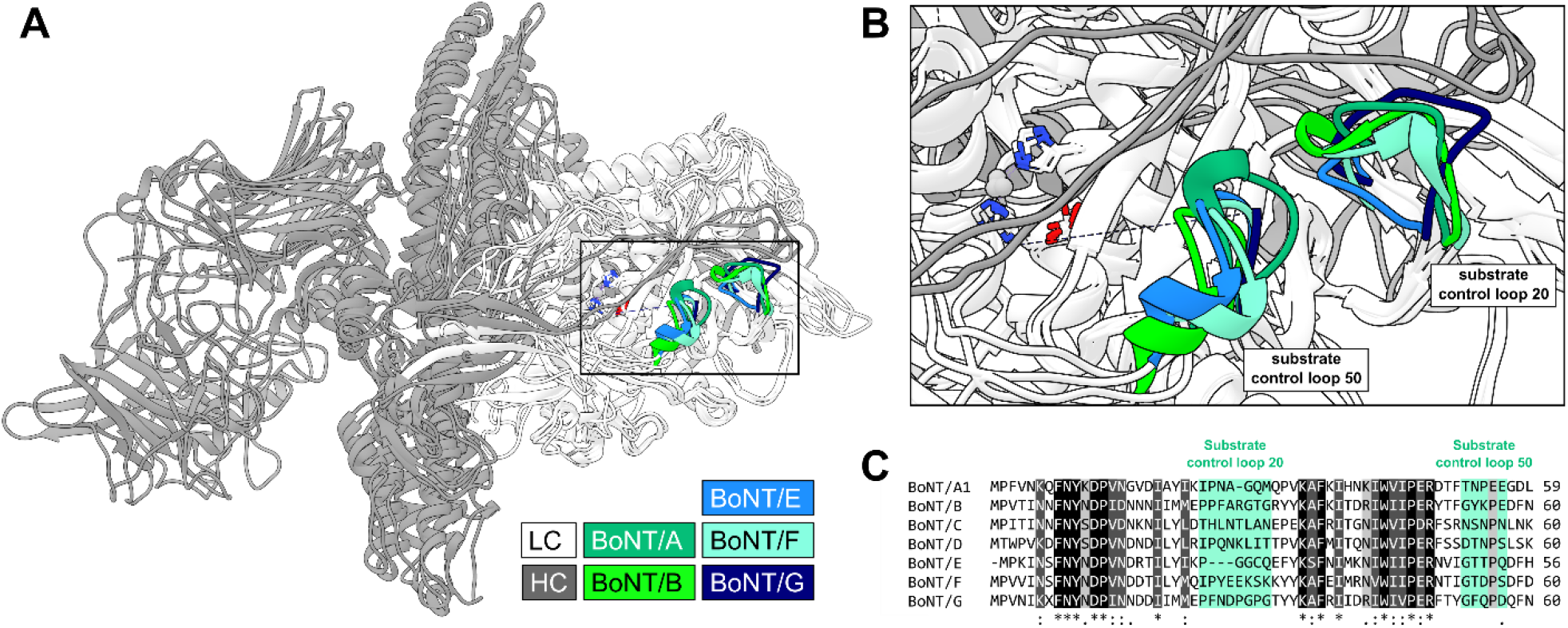
Substrate control loops are structurally present in other BoNT LC serotypes. (**A**) Multiple BoNT holotoxins were structurally aligned with UCSF Chimera MatchMaker (PDB IDs: 3BTA, 1S0G, 3FFZ, 1ZB7, & 2A8A). The heavy chain (HC) of each BoNT is colored dark grey, the light chain (LC) is colored in white, and the substrate control loops are colored as indicated in the key. (**B**) Larger image of the boxed section in A highlighting the substrate control loops. (**C**) The primary sequences of LC serotypes A through G were aligned using Clustal Omega. The substrate control loops are highlighted and annotated in teal. Identical residues are color-coded black (*), strongly similar residues in dark grey (:), weakly similar residues in light grey (.), and dissimilar residues are transparent. The poor primary sequence conservation in these structurally conserved loops suggests they help inform the diverse substrate specificities of other LC serotypes.

